# Identification of biomarkers for ulcerative morphology in carotid atherosclerosis plaques through single-cell RNA sequencing data

**DOI:** 10.1101/2025.07.14.664831

**Authors:** Huai Wu Yuan, Boyan Song, Hongzhe Wang, Tian Xiang Chen, Yun Zhen Hu, Guo Ping Peng

**Affiliations:** Department of neurology, the first affiliated hospital of Zhejiang university school of medicine, No. 1367 Wenyi West Road, Hangzhou City, Zhejiang Province, China; Lianchuan Biotechnology Co., Ltd., No. 758 Weiken Street, Hangzhou City, Zhejiang Province, China; Department of Clinical Pharmacy, the first affiliated hospital of Zhejiang university school of medicine, No. 1367 Wenyi West Road, Hangzhou City, Zhejiang Province, China

**Keywords:** Ulcerative plaques, Carotid atherosclerosis, Single-cell RNA sequencing, Biomarkers

## Abstract

**Background:** Carotid atherosclerotic stenosis (CAS) is a major risk factor for stroke, particularly when associated with ulcerative plaque morphology. However, the molecular mechanisms underlying ulceration remain inadequately defined. In this study, we aimed to construct single-cell atlases of ulcerative and non-ulcerative CAS plaques to identify biomarkers associated with ulcerative lesions.

**Methods:** Plaque tissues from four ulcerative and three non-ulcerative CAS samples were collected and subjected to single-cell RNA sequencing (scRNA-seq). Cell types were identified through scRNA-seq analysis, and differential cell types were determined by comparing their proportions between the 2 groups. Candidate genes were identified by intersecting differentially expressed genes (DEGs) within these differential cell populations. Subsequently, the Molecular Complex Detection (MCODE) plugin in Cytoscape was employed to confirm the biomarkers. Functional annotation and regulatory network analysis were then performed to investigate underlying biological mechanisms.

**Results:** scRNA-seq analysis identified 7 cell types, of which 6 showed differential proportions, leading to the identification of 61 candidate genes and 7 key biomarkers (KLF2, JUNB, FOS, HSPA1A, DUSP1, JUND, and ZFP36). Enrichment analysis indicated that these biomarkers may influence ulcerative CAS plaques via RNA polymerase II-mediated transcription and stress-response pathways. Regulatory network analysis revealed complex interactions, with RBMX, STAT3, YTHDF3, DDX3X, and ELAVL1 identified as potential upstream regulators.

**Conclusions:** KLF2, JUNB, FOS, HSPA1A, DUSP1, JUND, and ZFP36 were identified as biomarkers, offering novel insights into the molecular mechanisms underlying ulcerative morphology in CAS.

## Introduction

Ulceration is a key morphological feature of advanced carotid artery plaques, and is associated with increased stroke risk.^1–2^ Intraplaque hemorrhage (IPH), a hallmark of plaque vulnerability, is frequently observed in ulcerated plaques and may contribute to rupture by promoting inflammation and neovessel leakage.^3–4^ Elevated shear stress on the plaque surface may also trigger rupture, subsequently resulting in ulceration.^5–7^ However, ulceration is often underdiagnosed by conventional imaging (e.g., ultrasound or CTA) due to its microscopic size or superficial morphology.^8^ Notably, ulcerated plaques may remain asymptomatic without causing overt cerebrovascular events, highlighting the complexity of their pathogenic mechanisms.^9^

Bulk RNA-seq reveals transcriptomic profiles across entire tissue samples but obscures among individual cells. Single-cell RNA sequencing (scRNA-seq) addresses this limitation by enabling high-resolution mapping of cell populations and their biological characteristics. scRNA-seq offers key advantages, including identification of novel cell types, monitoring of dynamic cellular changes, analysis of cellular subpopulations, and assessment of heterogeneity at the single-cell level. These strengths enable more detailed and accurate disease investigations.^10–12^

This study utilized scRNA-seq data from ulcerated and non-ulcerated carotid artery plaque tissues in CAS, combined with bioinformatics approaches to investigate the cellular atlas of both sample groups. We systematically analyzed cell populations differing in abundance between the groups and subsequently identified biomarkers specific to ulcerated CAS plaques. Building on these biomarkers, we conducted multi-dimensional analyses, including functional annotation, regulatory network construction, drug prediction, and molecular docking simulations to elucidate their biological relevance.

## Methods

### Data Availability Statement

This study was approved by the Ethics Committee of the First Affiliated Hospital of Zhejiang University (IIT20241520A) and was conducted by the principles of the Declaration of Helsinki. Written informed consent was obtained from all patients or their legal representatives.

### Sample collection and scRNA-seq

In this study, we collected tissue samples from four and three ulcerative (case group) and non-ulcerative plaques (control group) from patients with CAS to prepare single-cell suspensions for scRNA-seq. During the carotid artery stenting procedure, the balloon was inflated to compress the ulcerated plaque, and dislodged emboli were captured using an embolic protection system. Following the procedure, the balloon and protection system were excised, placed in cell preservation solution, and immediately transported to the laboratory for further processing. After cell lysis across groups, additional dilution and separation steps were performed to generate single-cell suspensions. Subsequently, 10xionthat the cells were reduced to index transcripts for each cell, employing approximately 750,000 unique barcodes. This process involved encapsulating thousands of cells into nanoliter-sized Gel Bead-in-Emulsion droplets. The individual cells were then lysed to release RNA for downstream library preparation. Finally, scRNA-seq was performed using Illumina sequencing platforms (Illumina, San Diego, CA, USA).

### The scRNA-seq analysis

To explore the cellular mechanisms underlying case and control samples, a series of analyses were conducted on the scRNA-seq data using the Seurat (v 5.1.0) package.^13^ Initial steps involved quality control (QC) measures to remove low-quality data potentially resulting from cell damage or library preparation defects, implemented via the PercentageFeatureSet function in Seurat (v 5.1.0). QC was performed using the following criteria: (1) genes expressed in fewer than three cells were excluded, (2) cells with <200 or >5,000 genes were excluded, (3) cells with total gene expression counts >10,000 were excluded, and (4) cells with >5% mitochondrial gene expression were excluded.

Following QC, data normalization of the retained samples was performed using the NormalizeData function, applying a scale factor of 10,000. Highly variable genes (HVGs) were identified using the FindVariableFeatures function, and the LabelPoints function was used to visualize the top 10 genes with the most marked expression variability. To assess cell distribution uniformity across samples and identify potential outliers, the ScaleData function was applied. Principal component analysis (PCA) was then conducted using the selected HVGs to reduce data dimensionality. The top 50 principal components (PCs) were calculated to extract the most informative variance features. To determine the optimal number of PCs, the JackStraw method was employed to assess the contribution of the top 50 PCs (p < 0.05), and results were visualized using the JackStrawPlot function. Simultaneously, the ElbowPlot function generated a scree plot to visualize the contribution of each PC.

Next, cell cluster analysis was performed on the data following dimensionality reduction using the FindNeighbors and FindClusters functions (resolution = 0.4). The resulting cell clusters were visualized using t-distributed stochastic neighbor embedding (t-SNE). Subsequently, cluster annotation was conducted based on marker genes identified via the FindAllMarkers function (minpct = 0.40, log2Fold Change (FC) = 0.25, p < 0.05) and cross-referenced with literature data.^14^ The annotated cell types were visualized with t-SNE, and the expression patterns of the corresponding marker genes within each cell type were further depicted using a bubble chart.

### Gene set variation analysis (GSVA) and recognition of candidate genes

To explore pathway differences between case and control groups, the GSVA (v 1.50.0) package was used to perform GSVA.^15^ The analysis applied the “c2.cp.kegg.v7.4.symbols.gmt” background gene set from the Molecular Signatures Database (MSigDB) (https://www.gsea-msigdb.org/gsea/msigdb). Enrichment scores for various pathways were obtained, and differences in functional enrichment between the groups were compared using the limma (v 3.54.0) package (|t| > 2, p < 0.05).^16^ Results were visualized in a bar chart.

Subsequently, the proportions of annotated cell types between the case and control samples were compared using the Fisher test, and cell types showing significant proportional differences between the two groups were defined as differential cell types (p < 0.05). Additionally, cell cycle state analysis of the differential cell types was performed using the scran (v 1.30.0) package.^17^ Differentially expressed genes (DEGs) within each differential cell type between the case and control samples were then identified using the FindMarkers function (|average log_2_FC| > 0.1, p < 0.05). Volcano plots were generated to visualize the DEGs in each differential cell type using the ggplot2 (v 3.5.1) package, with the top 10 upregulated and downregulated genes labeled based on |average log_2_FC| (ranked from high to low).^18^ Further analysis involved identifying the intersection of DEGs across differential cell types to obtain candidate genes, which were visualized using the ggVennDiagram (v 1.5.2) package.^19^

### Enrichment analysis of candidate genes

To investigate the cellular functions of candidate genes and the pathways in which they were involved, Gene Ontology (GO) and Kyoto Encyclopedia of Genes and Genomes (KEGG) enrichment analyses were performed using the clusterProfiler (v 4.10.1) package (p < 0.05).^20^ GO analysis encompasses three main categories: biological process (BP), cellular component (CC), and molecular function (MF). Results were ranked in descending order based on gene count, and the top 5 terms for BP, CC, and MF, along with the top 5 significant KEGG pathways, were selected for presentation.

### Recognition of biomarkers

After identifying the candidate genes, these genes were uploaded to the Search Tool for the Retrieval of Interacting Genes/Proteins (STRING) database (http://string-db.org) to obtain their protein-level interactions (confidence > 0.4). These interactions were visualized using Cytoscape (v 3.9.1) software to construct a protein-protein interaction (PPI) network.^21^ In addition, the Molecular Complex Detection (MCODE) plugin in Cytoscape (v 3.9.1) was employed to identify key modules (Degree Cutoff = 2, Node Score Cutoff = 0.2, K-Core = 2, and Max Depth = 100), with resulting interactions also visualized using Cytoscape (v 3.9.1). The genes in these key modules were subsequently analyzed using the GeneCards database (https://www.genecards.org/) to retrieve their functional annotations. The key module containing genes involved in cell proliferation, differentiation, and death was identified, and the genes within this module were selected as biomarkers for subsequent analysis.

### Comprehensive functional characterization analysis of biomarkers

After identifying the biomarkers, a series of analyses were conducted to explore their characteristics. First, Spearman correlation analysis was performed among the biomarkers (|correlation coefficient (cor)| > 0.30, p < 0.05) using the psych (v 2.4.3) package.^22^ Next, functional similarity scores (> 0.50) among the biomarkers were calculated using the GOSemSim (v 2.26.1) package to examine their functional relationships.^23^ Additionally, to investigate their chromosomal localization, the positions of the biomarkers on chromosomes were visualized using the OmicCircos (v 1.40.0) package.^24^ Finally, genes associated with the functions and shared activities of the biomarkers were predicted using the GeneMANIA database (http://www.genemania.org/), leading to the construction of a gene-gene interaction (GGI) network for visualization.

### Construction of regulatory networks

To delve into the molecular regulatory mechanisms governing these biomarkers, relationships between transcription factors (TFs) and biomarkers were predicted using the Transcriptional Regulatory Relationships Unraveled by Sentence-based Text mining (TRRUST) database (https://www.grnpedia.org/trrust/), enabling the construction of a TF-mRNA regulatory network. Subsequently, to identify RNA-binding proteins (RBPs) associated with these biomarkers, a comprehensive search was conducted using the Starbase database (http://starbase.sysu.edu.cn/). An interaction network was then constructed to map the connections between RBPs and biomarkers. The resulting regulatory networks were graphically represented using Cytoscape (v 3.9.1) software.

### Heterogeneity of endothelial cells and analysis of cell communication

To gain deeper insights into the heterogeneity of endothelial cells, the endothelial cells were divided into different subclusters by performing dimensionality reduction and clustering analysis, with the FindNeighbors and FindClusters functions (resolution = 0.2). These subclusters were further visualized using t-SNE. To decipher the molecular dialogues and infer the intricate interactions between annotated cell types, the CellChat (v 1.6.1) package was engaged for communication analysis within the annotated cell types, focusing on the endothelial cells.^25^ The study also presented a comprehensive visualization of receptor-ligand interplays through heatmaps (p < 0.05), depicting the exchange of signals received and dispatched by the annotated cell types, emphasizing the endothelial cells.

### Pseudo-time analysis

Subsequently, in order to investigate the differentiation trajectory and evolution of different subclusters in endothelial cells during development, the monocle (v 2.30.1) package was employed to conduct pseudo-time series analysis.^26^ Additionally, the plot_genes_in_pseudotime function was used to visualize the expression profiles of biomarkers across different differentiation states of endothelial cells, as well as between the case and control samples.

### Statistical analysis

All analyses were executed in R (v 4.2.2) software. To determine whether there were statistical differences between the two groups, the Fisher test was employed. A p-value < 0.05 was considered statistically significant.

## Results

### Annotation of seven cell types and exploration of biological pathway differences between case and control samples

scRNA-seq analysis was performed to explore the underlying mechanisms in case and control samples at the cellular level. Initially, visualizations of nFeature RNA (genes measured per cell), nCount RNA (total gene expression counts per cell), and percent mitochondrial content before and after quality control were generated (Supplementary Figures 1A–1B). A total of 38,114 cells and 34,270 genes were retained after filtering. Following standard preprocessing, 2,000 HVGs were selected for further analysis, and the top 10 most variable genes were highlighted (Supplementary Figure 1C). PCA showed no batch effect between the groups (Supplementary Figure 1D), and the top 30 PCs were selected for downstream analysis (Fig. 1A, Fig. 1B). Clustering analysis revealed 18 distinct cell clusters (Fig. 1C), which were annotated into seven cell types: B cells, CD8+T cells, CD4+T cells, mast cells, endothelial cells, monocytes, and neutrophils (Fig. 1D, Fig. 1E). A bubble chart demonstrated strong marker gene specificity, confirming the accuracy of cell-type annotations (Fig. 1F).

**Figure 1.**
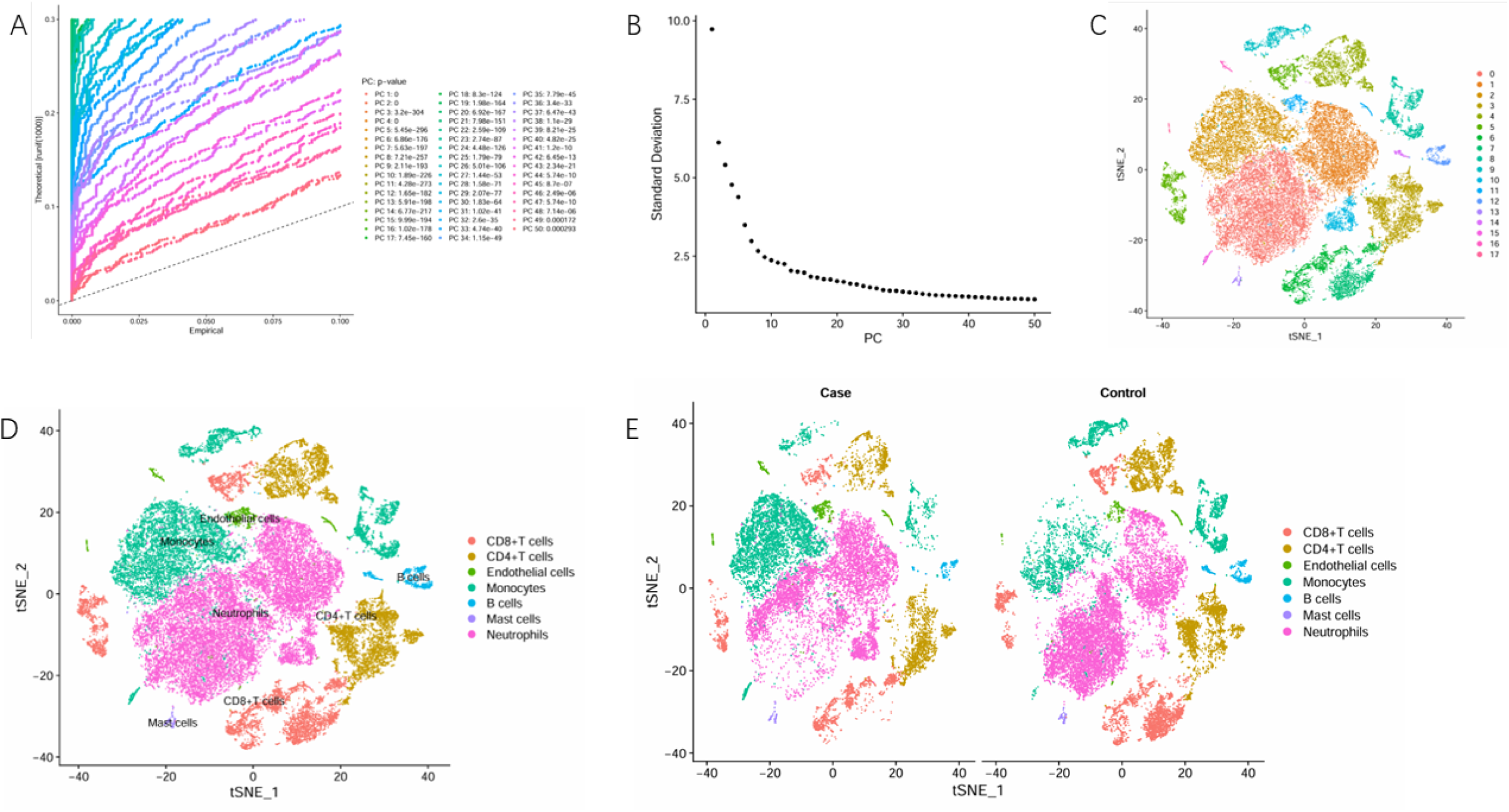
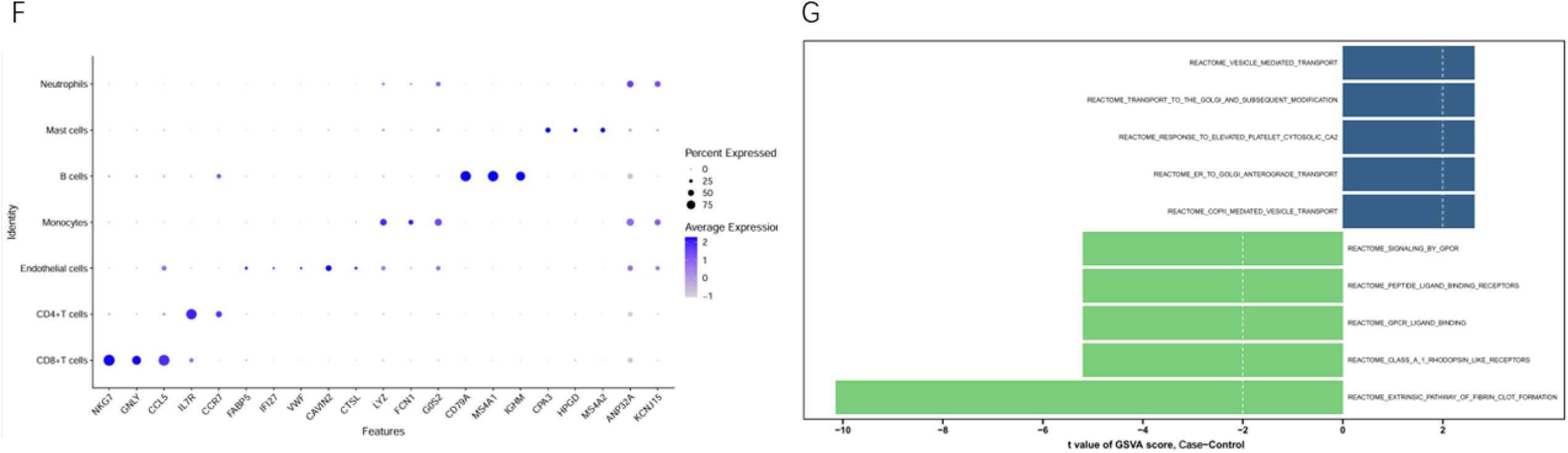
JackStraw plot (A) and identification of available dimensions (B) (In panel A, the x-axis represents experimental values, the y-axis represents theoretical values, and the points on the right indicate p-values. In panel B, the x-axis shows the number of principal components, and the y-axis displays the standard deviation. t-SNE algorithm-based dimensionality reduction and clustering visualization of cell subpopulations (C). Cell annotation results (D). Comparison of t-SNE between the case group and the control group (E). Bubble plot showing gene expression across different cell types (F). GSVA enrichment analysis of case and control samples. (The x-axis represents t-values, with blue indicating pathways having t-values >2 and green indicating pathways with t-values < -2) (G).

Following the GSVA conducted between the case and control groups, 30 significant differential pathways were identified (|t| > 2, p < 0.05), including 11 activated and 19 suppressed pathways in the case samples (Supplementary Table 1). Notably, pathways such as “vesicle-mediated transport,” “response to elevated platelet cytosolic Ca²+,” and “COPII-mediated vesicle transport” were predominantly activated in case samples, indicating an elevated demand for protein processing and intracellular trafficking (Fig. 1G). In contrast, pathways such as “G-protein coupled receptor (GPCR) ligand binding,” “peptide ligand binding receptors,” and “signaling by GPCR” were suppressed, possibly reflecting dysregulation in fundamental signaling processes (Fig. 1G). These findings help elucidate the complex molecular mechanisms linked to cellular stress responses, protein processing dysfunction, and signaling irregularities, which may play a central role in the pathogenesis and progression of ulcerative plaques in CAS.

### Recognition and enrichment analysis of 61 candidate genes

Subsequently, the proportions of cell types were compared between the case and control samples, revealing 6 differential cell types: CD8+ T cells, CD4+ T cells, endothelial cells, monocytes, B cells, and neutrophils (p < 0.05) (Fig. 2A). Specifically, CD8+ T cells, CD4+ T cells, B cells, and neutrophils were significantly more abundant in the control group, whereas endothelial cells and monocytes were significantly prevalent in the case group. Cell cycle analysis showed that in both the G1 and G2M phases, neutrophils had the highest proportion, followed by monocytes (Fig. 2B). In the S phase, CD8+ T cells and CD4+ T cells exhibited higher proportions (Fig. 2B). These findings suggest that these immune and vascular cell types play distinct roles across cell cycle phases, offering insights into their contributions to homeostasis or disease progression.

**Figure 2.**
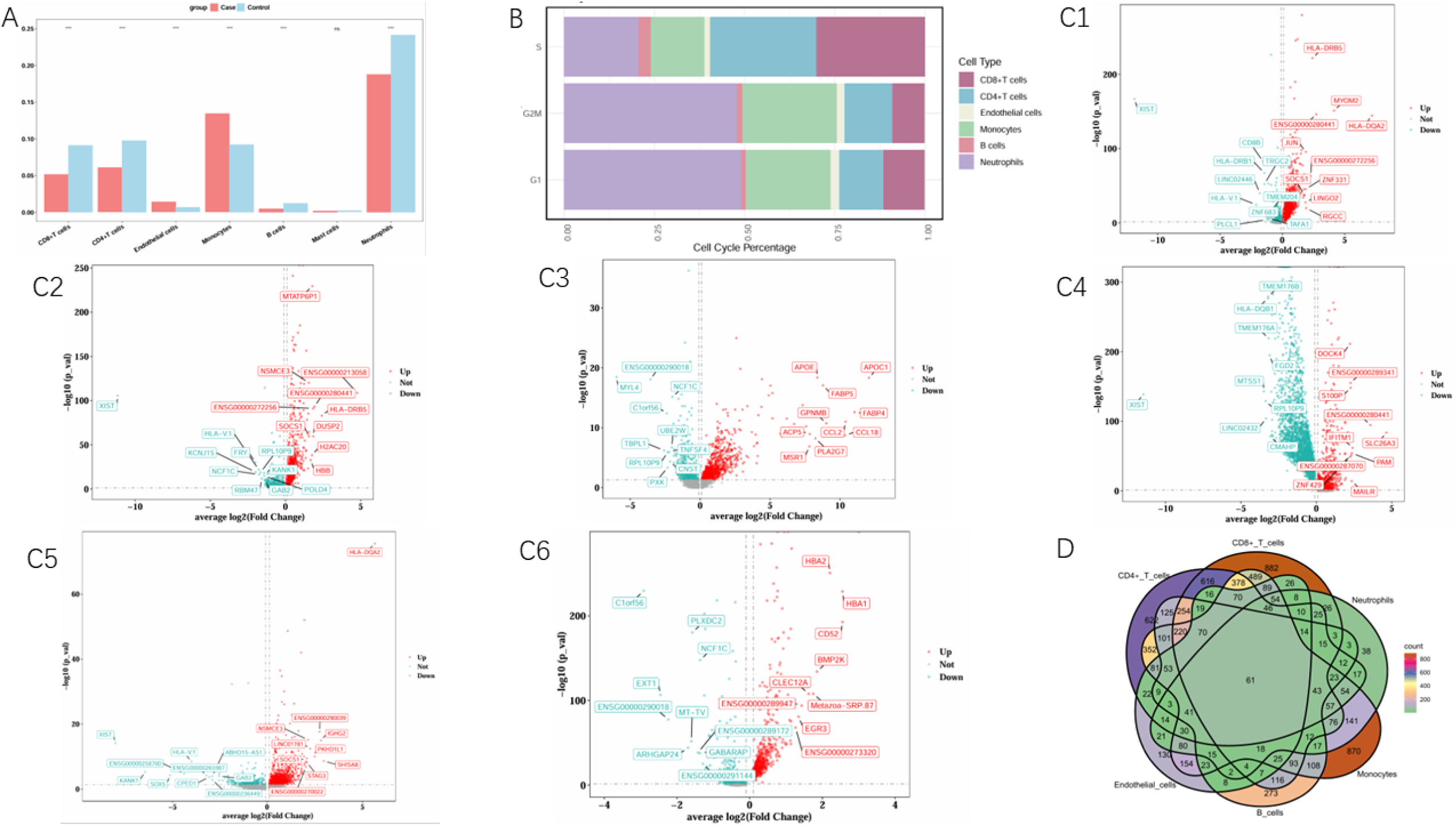
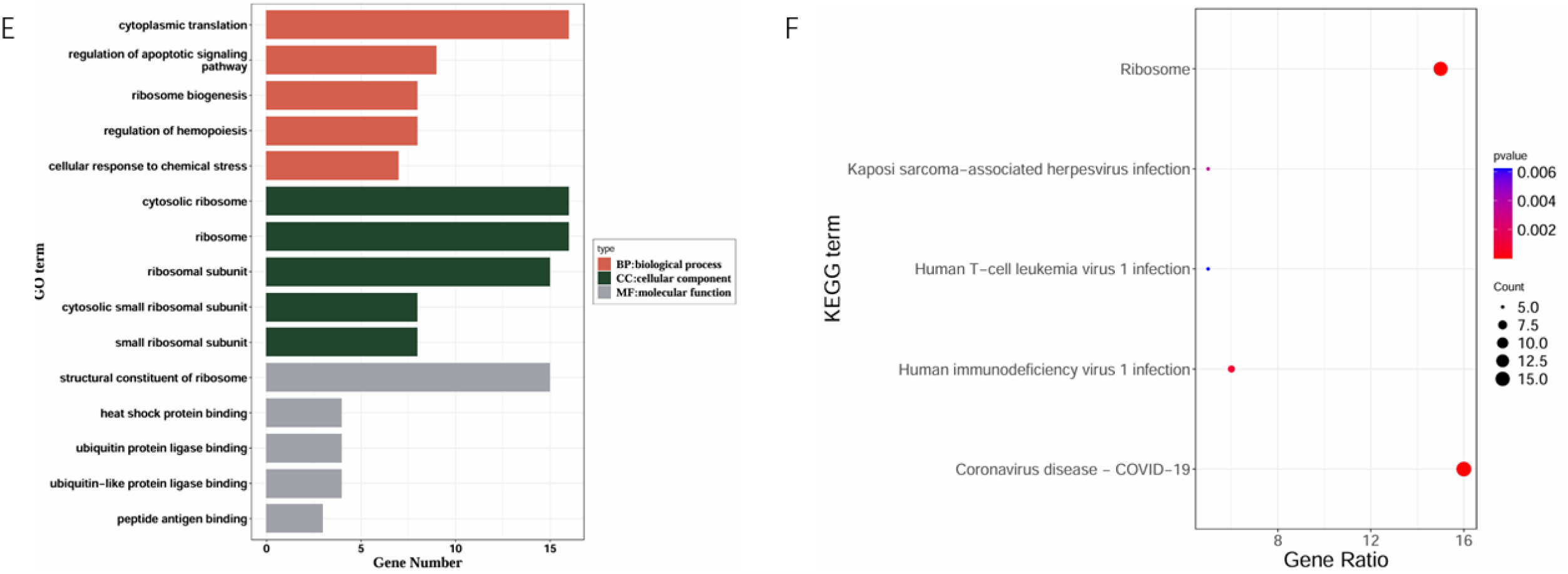
Cell types with percentage differences (A) (Red represents the CAS group, blue represents the control group). Cell cycle analysis of differential cells (B) (X-axis shows cell proportion, Y-axis shows cell cycle phase, with different colors indicating distinct cell types). Distribution of differentially expressed genes (DEGs) in differential cells between disease and normal control groups (C1-C6) (C1, DEGs in CD8+ T cells; C2, DEGs in CD4+ T cells; C3, DEGs in endothelial cells; C4, DEGs in monocytes; C5, DEGs in B cells; C6, DEGs in neutrophils). Intersection plot of DEGs from differential cells (D) (Colors represent gene counts, with darker shades indicating higher numbers of genes). GO enrichment analysis of candidate genes (E) (Red represents Biological Processes [BP], green represents Cellular Components [CC], and gray represents Molecular Functions [MF]. The x-axis shows the number of enriched genes, with longer bars indicating higher gene counts. KEGG enrichment analysis of candidate genes (F) (The x-axis displays the number of enriched genes, where larger circles represent higher gene counts. The y-axis shows enriched pathways, with more intense red coloration indicating greater statistical significance.

Furthermore, differential expression analysis conducted on each differential cell type between the case and control groups revealed 4,566 DEGs in CD8+ T cells, 4,921 DEGs in CD4+ T cells, 2,341 DEGs in endothelial cells, 4,236 DEGs in monocytes, 5,236 DEGs in B cells, and 1,262 DEGs in neutrophils (|average log_2_FC| > 0.1, p < 0.05) (Fig. 2C, Supplementary Table 2). The intersection of these DEGs from each differential cell type led to the identification of 61 candidate genes (Fig. 2D, Supplementary Table 3), which may provide a foundation for investigating their functional roles in disease pathogenesis.

Further analysis of these 61 candidate genes using GO analysis identified 501 significant terms (p < 0.05), including 384 BP terms, 70 CC terms, and 47 MF terms (Supplementary Table 4). At the GO level, these candidate genes were primarily enriched in functions such as “cytoplasmic translation” (BP), “cytosolic ribosome” (CC), and “structural constituent of ribosome” (MF) (p < 0.05) (Fig. 2E). Additionally, 27 KEGG pathways were identified, including “ribosome,” “human immunodeficiency virus 1 infection,” and “Kaposi sarcoma-associated herpesvirus infection” (p < 0.05) (Fig. 2F, Supplementary Table 4). These findings underscore the potential involvement of these candidate genes in essential biological processes and signaling pathways, offering insights into their functional significance and contribution to disease mechanisms.

### Selection of KLF2, JUNB, FOS, HSPA1A, DUSP1, JUND, and ZFP36 as biomarkers

Following this, after removing 15 discrete proteins, the remaining candidate genes were visualized in the PPI network, with RPS3 and RPS2 exhibiting strong interactions with other genes (Fig. 3A). Moreover, the MCODE algorithm was applied, resulting in three key modules in the PPI network (Fig. 3B). Gene function retrieval of genes within these key modules was performed using GeneCards, revealing that genes in key module 1 were predominantly associated with ribosomal protein functions; genes in key module 2 were primarily involved in regulating cell proliferation, differentiation, and apoptosis; and genes in key module 3 were mainly linked to immune responses. Consequently, genes in key module 2 were selected as biomarkers for this study, namely KLF2, JUNB, FOS, HSPA1A, DUSP1, JUND, and ZFP36. These findings suggest that these biomarkers might play key roles in the development and pathogenesis of ulcerative plaques in CAS, highlighting their potential as therapeutic targets for treatment.

**Figure 3.**
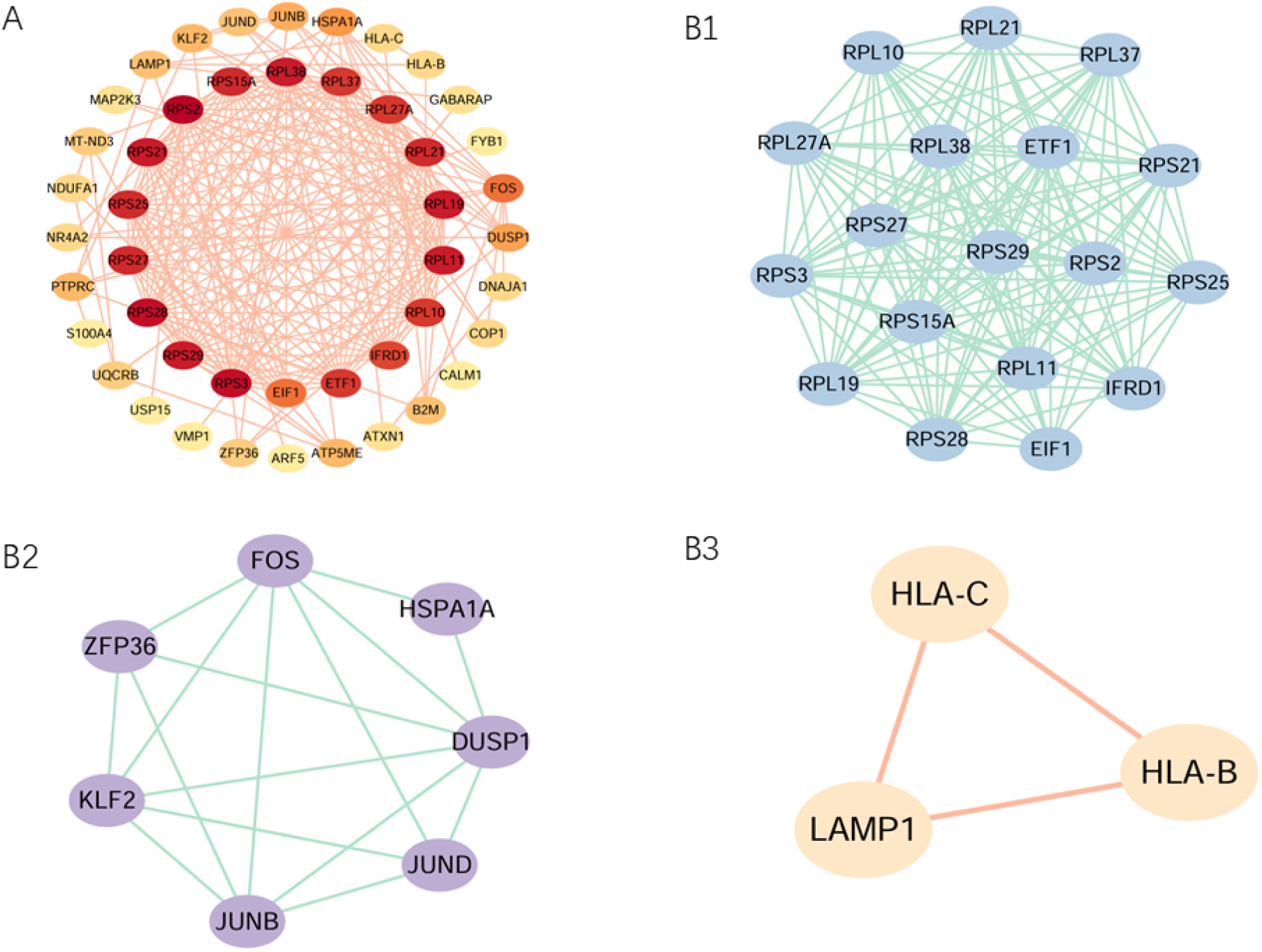
PPI network (A) (Node color intensity indicates connectivity strength, with redder hues representing higher degrees of interaction. Edges represent gene-gene interactions). MCODE subnetworks (B1-B3 correspond to Module 1, Module 2, and Module 3, respectively).

### Functional analyses of seven biomarkers

Based on the Spearman correlation analysis, most biomarkers demonstrated significant positive correlations, with the strongest observed between DUSP1 and FOS (cor = 0.68, p < 0.05) (Fig. 4A, Supplementary Table 5). Except for DUSP1, all other biomarkers had scores >0.5, with JUNB showing the highest score, suggesting strong functional similarities among these biomarkers (Fig. 4B, Supplementary Table 6). Chromosomal localization analysis revealed that KLF2, JUNB, JUND, and ZFP36 were positioned on chromosome 19; FOS on chromosome 14; HSPA1A on chromosome 6; and DUSP1 on chromosome 5 (Fig. 4C). A GGI network was subsequently constructed using GeneMANIA, which identified 20 genes associated with the biomarkers, including JUN, FOSB, and ATF3. These genes were enriched in functions such as “RNA polymerase II cis-regulatory region sequence-specific DNA binding,” “positive regulation of transcription by RNA polymerase II,” and “RNA polymerase II-specific DNA-binding transcription factor binding” (Fig. 4D). Collectively, these findings suggest that the biomarkers and their interacting genes may contribute to complex regulatory mechanisms involved in transcriptional control.

**Figure 4.**
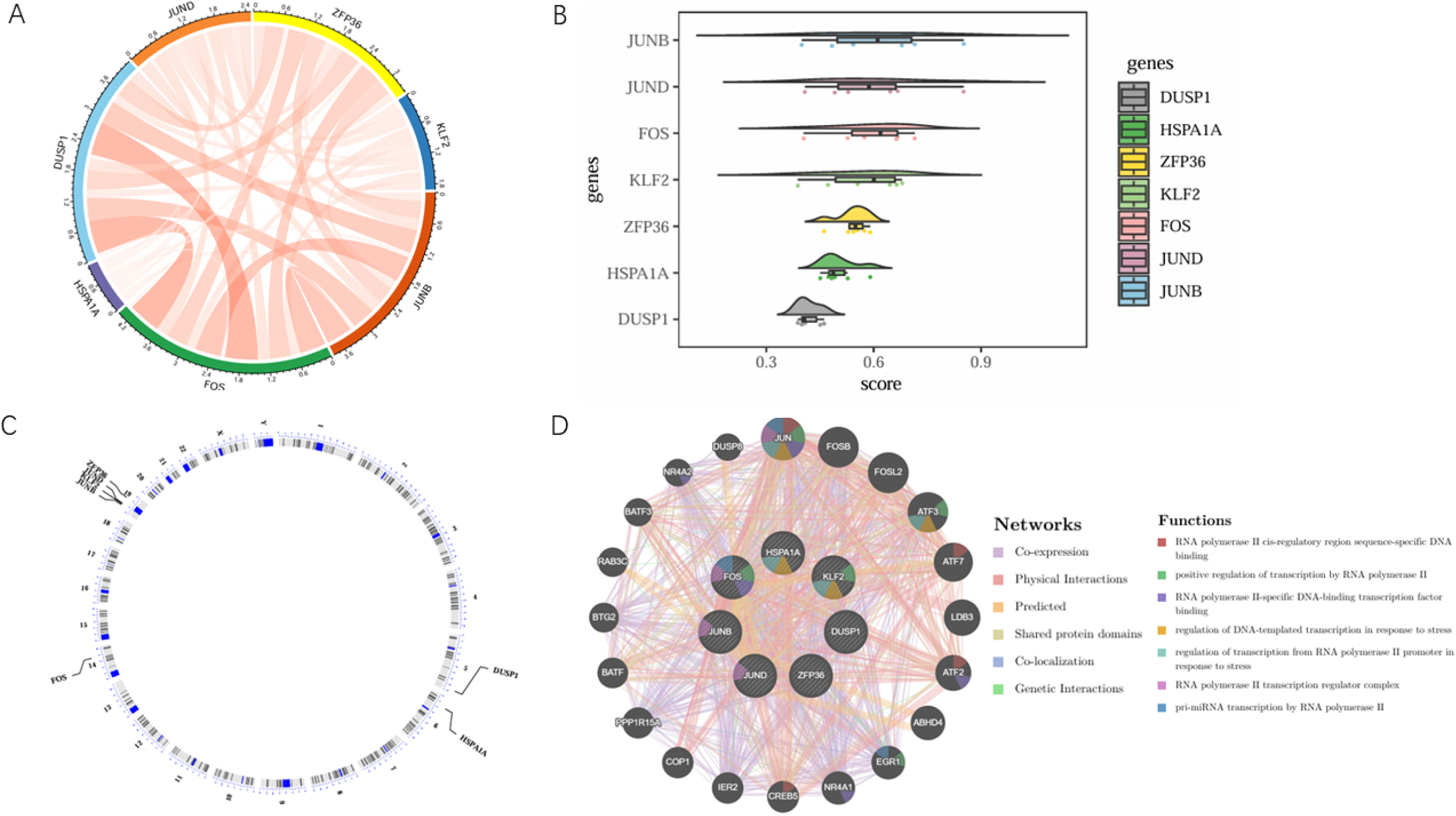
Correlation analysis of biomarkers (A) (Outer ring colors represent different genes, while inner colors indicate correlation strength—redder hues and thicker connecting lines denote stronger positive correlations). Functional similarity cloud-rain plot of biomarkers (B) (X-axis displays score values, Y-axis shows genes). Chromosomal mapping of biomarkers (C). GeneMANIA analysis (D) (Central large circle represents biomarkers, peripheral small circles indicate biomarker-associated genes. Network connections from top to bottom represent: co-expression, physical interactions, predictions, shared protein domains, colocalization, and genetic interactions. Functional annotations include: RNA polymerase II cis-regulatory region sequence-specific DNA binding, positive regulation of RNA polymerase II transcription, RNA polymerase II-specific DNA-binding transcription factor binding, DNA-templated transcription regulation in stress response, RNA polymerase II promoter transcriptional regulation, response to stress, RNA polymerase II transcription regulator complex, and pri-miRNA transcription via RNA polymerase II).

### Revealing regulatory networks for biomarkers

The TRRUST database predicted TFs regulating the biomarkers, identifying 50 TFs after merging and removing duplicates (4 for KLF2, 7 for JUNB, 31 for FOS, 3 for HSPA1A, 7 for DUSP1, 3 for JUND, and 4 for ZFP36) (Supplementary Table 7). These TFs were used to construct a TF-mRNA network (Supplementary Figure 2A). Within this network, certain TFs, such as RBMX and STAT3, may act as potential regulators due to their capacity to concurrently modulate multiple biomarkers. Subsequently, a total of 153 RBPs were predicted for the biomarkers using the StarBase database (Supplementary Table 8). Among these RBPs, 15 were associated with KLF2, 71 with JUNB, 38 with FOS, 3 with HSPA1A, 76 with DUSP1, 137 with JUND, and 52 with ZFP36. Based on these interactions, an RBP-mRNA regulatory network was constructed, highlighting potential regulators such as YTHDF3, DDX3X, and ELAVL1 (Supplementary Figure 2B). These findings provide a basis for further exploration of the functional roles of these regulatory interactions.

### Specific communications of cell types

The analysis of cellular communications among the 7 cell types further elucidated the extent and pathways of interactions between these cell types, clearly revealing significant differences in cell communication patterns between case and control samples. Compared to the control sample, case samples exhibited increased numbers and weights of cellular communications, indicating more frequent and functionally robust interactions among these cell types (Fig. 5A, Fig. 5B). Specifically, in case samples, endothelial cells exhibited the highest number of interaction with monocytes and neutrophils (Fig. 5C, Fig. 5D). Additionally, the strongest interaction weights were observed between endothelial cells and monocytes in case samples, as well as between endothelial cells and neutrophils (Fig. 5C, Fig. 5D).

**Figure 5:**
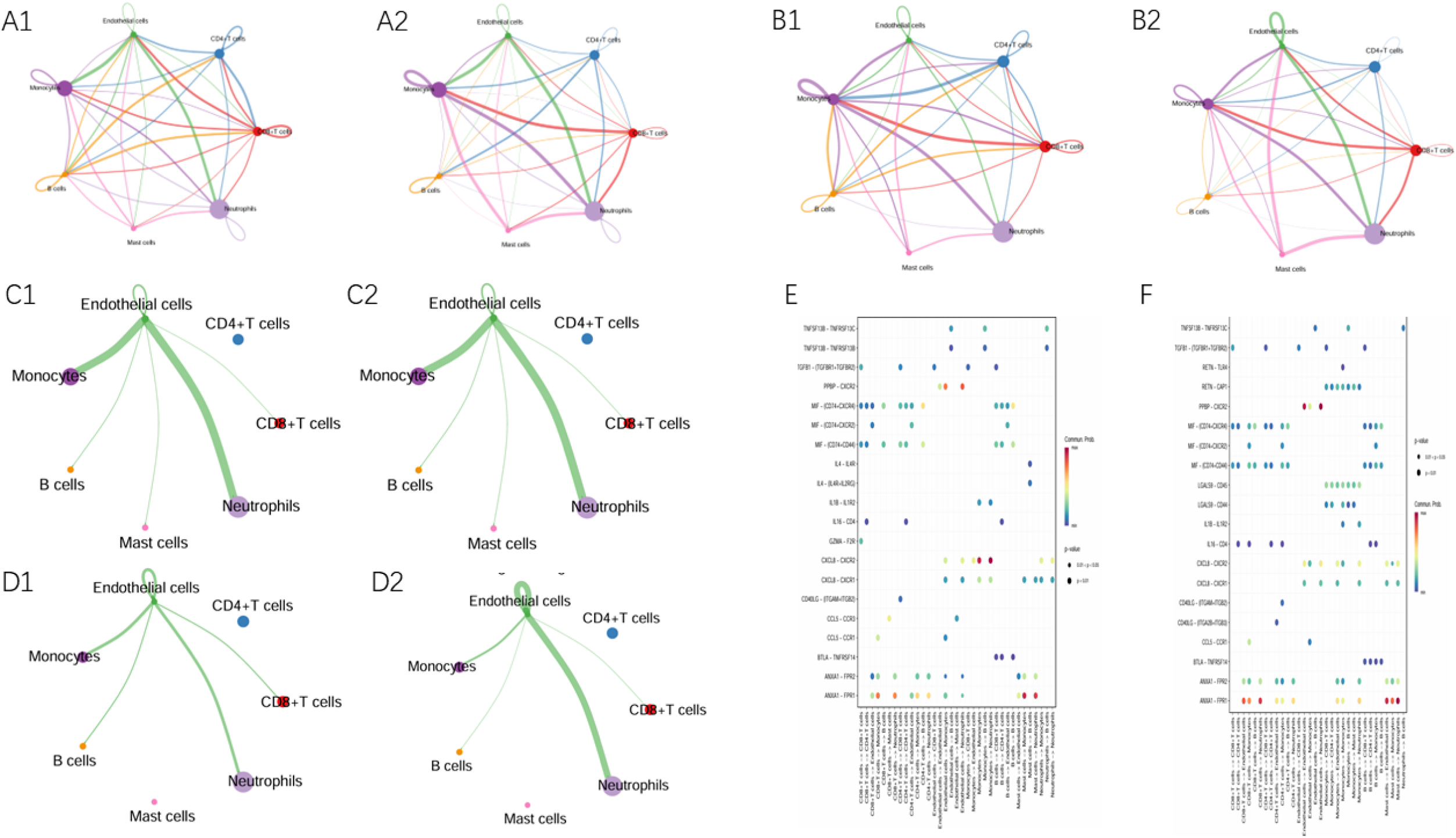
Cell communication analysis A1, case_net_number; A2, case_net_weight; B1, control_net_number; B2, control_net_weight; C1, case_number_Endothelial cells; C2, case_weight_Endothelial cells; D1, control_number_Endothelial cells; D2, control_weight_Endothelial cells; E, case dotplot; F, control dotplot.

Additionally, in case samples, enhanced communication was observed between endothelial cells and monocytes, as well as between endothelial cells and neutrophils, with the PPBP-CXCR2 ligand-receptor interaction playing a pivotal role (Fig. 5E). In control samples, a pronounced interaction was noted between endothelial cells and neutrophils, also involving the PPBP-CXCR2 ligand-receptor interaction (Fig. 5F). Future studies could focus on functional assays to better understand the significance of these interactions.

### Pseudo-time trajectory inference of endothelial cells

The following investigation was dedicated to endothelial cells. After dimensionality reducing and clustering analysis, endothelial cells were divided into 4 subclusters (Fig. 6A, Fig. 6B). Subsequently, the trajectories of endothelial cells were inferred by cell trajectory analysis and categorized into 3 parts (Fig. 6C). The starting point of the proposed temporal trajectory of endothelial cells was determined, indicating that as the cells progressed away from this starting point, they underwent maturation in their development. The intensity of the color in the graph was indicative of the timing of cell differentiation, with darker shades representing earlier periods. The subcluster 0 was more highly expressed in state 1, the subcluster 3 was more highly expressed in state 2, and the subcluster 2 was more highly expressed in state 3 (Fig. 6C). The case and control samples were found in all states (Fig. 6C).

**Figure 6:**
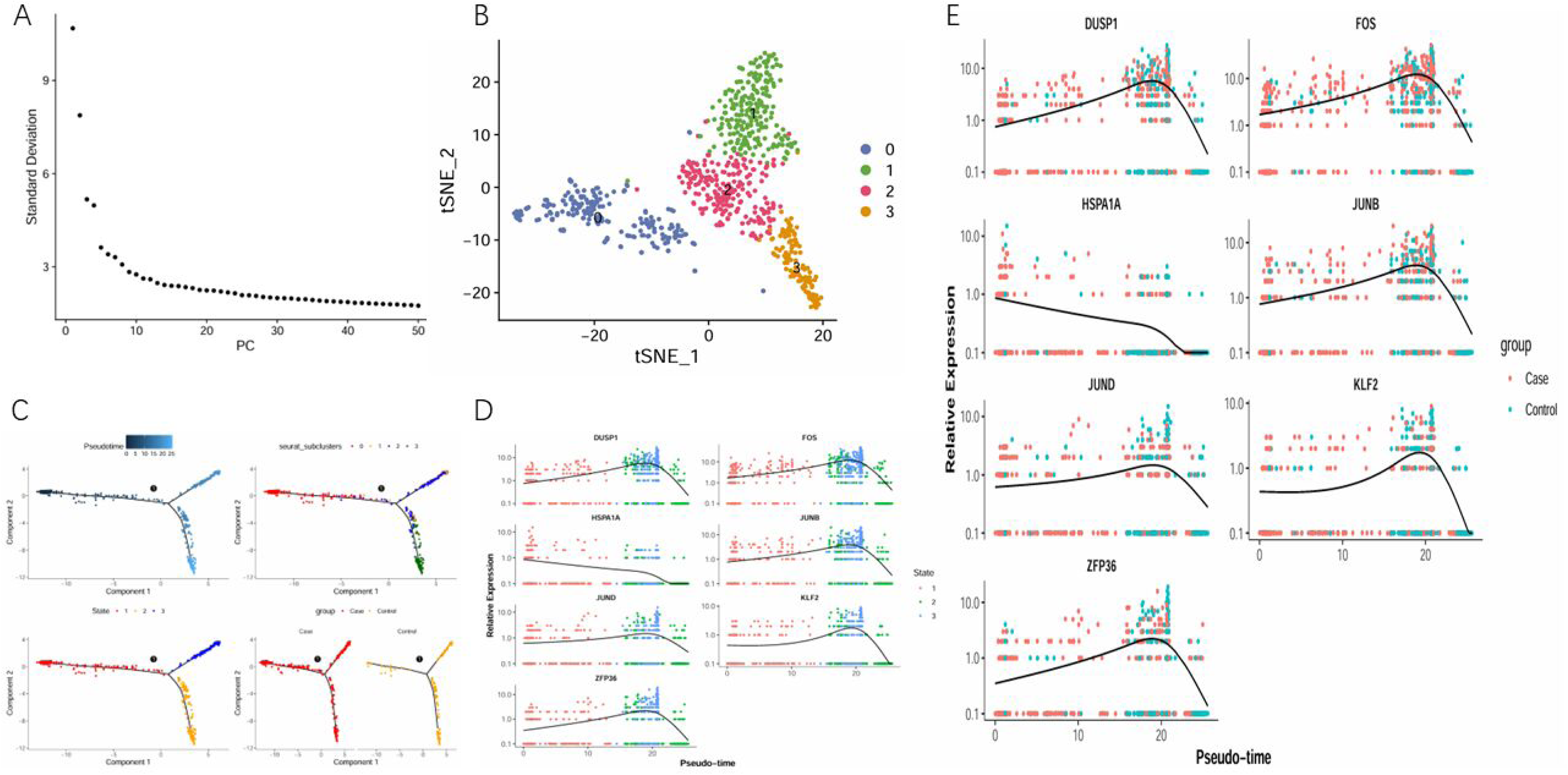
Pseudo-time trajectory inference of endothelial cells A, Endothelial_subcluster_ElbowPlot; B, subcluster_Endothelial_tsne; C, Endothelial_monocle; D, Endothelial_State_gene_exp; E, Endothelial_group_gene_exp.

Furthermore, the expression levels of biomarkers were also observed along the pseudo-time trajectory across different differentiation states and groups (Fig. 6D, Fig. 6E). Specifically, the expression patterns of all biomarkers exhibited complexity. The expression levels of 6 biomarkers (KLF2, JUNB, FOS, DUSP1, JUND, and ZFP36) initially increased and then decreased over time, while the expression level of HSPA1A consistently decreased throughout the endothelial cell differentiation process. These findings suggested that the expression levels of these biomarkers were dynamically regulated during endothelial cell differentiation, potentially reflecting their distinct biological roles at different stages of differentiation.

## Discussion

In this study, we performed single-cell RNA sequencing to comparatively analyze ulcerated versus non-ulcerated carotid plaques, identifying seven key biomarkers (KLF2, JUNB, FOS, HSPA1A, DUSP1, JUND, and ZFP36) associated with ulcerative morphology. Functional analyses revealed their roles in transcriptional regulation and stress response pathways. Through regulatory network mapping, we further elucidated their mechanistic relationships and potential therapeutic targeting.This study identified cell types at the single-cell level through annotation (comprising 7 cell types: B cells, CD8+ T cells, CD4+ T cells, mast cells, endothelial cells, monocytes, and neutrophils). Subsequently, differential abundance analysis was performed to identify significantly altered cell types (six cell types: CD8+ T cells, CD4+ T cells, endothelial cells, monocytes, B cells, and neutrophils). Previous studies have shown that cardiovascular disease susceptibility genes are particularly enriched in lesional macrophages, endothelial cells, and smooth muscle cells.^27–28^ LYVE-1 res-like macrophage accumulation may be associated with plaque rupture in human carotid artery disease.^29^ Our findings further support that the monocyte–macrophage system and endothelial cells play important roles in the pathogenesis of ulcerative plaque features.

This study utilized single-cell sequencing data from ulcerated and non-ulcerated plaques in carotid atherosclerosis (CAS) and identified seven biomarkers through systematic analytical screening. KLF2 protects blood vessels from injury and slows atherosclerosis progression by suppressing pro-inflammatory factor expression in endothelial cells while promoting anti-inflammatory genes. Reduced KLF2 expression is closely linked to plaque formation and rupture, likely affecting plaque stability through its regulatory influence on endothelial function.^30–32^ JUNB, FOS, and JUND belong to the AP-1 transcription factor family. JUNB plays a key role in plaque formation, and its overexpression may contribute to rupture and instability.^33–34^ FOS is critical to atherosclerosis progression, regulating activation in endothelial, smooth muscle, and immune cells, whose overactivation contributes to plaque destabilization.^35–36^ Compared to other Jun family members, JUND may play distinct roles in cell proliferation, often associated with growth cycles and responses to external signals.^37–38^ HSPA1A (Heat shock protein family A member 1A) protects cells from stress-induced damage, regulates protein folding, and supports cellular stability. Genetic variation in HSPA1A may be linked to atherosclerosis.^39^ DUSP1 is a dual-specificity phosphatase regulating pathways such as MAPK; it may modulate inflammation and cellular stress in atherosclerosis.^40^ ZFP36, a zinc finger transcriptional repressor, regulates mRNA stability and pro-inflammatory cytokine expression, modulating immune responses.^41^ In summary, current evidence supports that KLF2 and AP-1 transcription factors may play active roles in the formation of ulcerative plaque features.

Except DUSP1, the functional similarity scores of the remaining biomarkers were all >0.5. This implies that these biomarkers may share overlapping functions or signaling pathways in the disease context and may collectively contribute to disease progression through synergistic mechanisms. Notably, ZFP36, KLF2, JUNB, and JUND are all located on chromosome 19, suggesting that they may engage in related biological processes. Their co-localization may support coordinated transcriptional regulation among these genes. Additionally, the biomarkers and their associated genes are predominantly involved in biological processes such as “RNA polymerase II cis-regulatory region sequence-specific DNA binding,” “positive regulation of transcription by RNA polymerase II,” and “RNA polymerase II-specific DNA-binding transcription factor binding.” These functional associations indicate that the biomarkers might influence ulcerative plaques in CAS by modulating RNA polymerase II-mediated transcription and stress-responsive pathway.^42–43^ Furthermore, this study suggests that RBMX, STAT3, YTHDF3, DDX3X, and ELAVL1 may be associated with multiple biomarkers. STAT3 can reportedly regulate inflammatory processes driving atherosclerotic plaque formation through diverse molecular mechanisms.^44–45^

This study’s in-depth analysis of endothelial cells reveals their critical role in ulcerated plaques of carotid atherosclerosis. Through secondary dimensionality reduction clustering, we identified four endothelial cell subpopulations, indicating significant heterogeneity within these cells in the plaque microenvironment. Cell communication analysis revealed a marked enhancement in PPBP (pro-platelet basic protein)-CXCR2 ligand-receptor interactions between endothelial cells and monocytes/neutrophils in ulcerated plaques. This finding aligns with previously reported mechanisms where CXCR2-mediated inflammatory cell recruitment promotes plaque instability.^46^

Pseudotime analysis further demonstrated dynamic expression patterns of seven biomarkers during endothelial cell differentiation: KLF2, JUNB, FOS, DUSP1, JUND, and ZFP36 were upregulated during the intermediate differentiation stage, potentially promoting endothelial functional maturation by regulating transcriptional activity and inflammatory responses.^47^ Conversely, HSPA1A expression showed a continuous decline, suggesting its role in maintaining endothelial homeostasis may diminish as differentiation progresses.

These dynamic features provide novel insights into the functional transformation of endothelial cells during plaque progression. Specifically, the coordinated upregulation of biomarkers during intermediate differentiation may promote plaque vulnerability by regulating extracellular matrix remodeling and inflammatory responses.^48^ Future studies should employ functional experiments to further validate the specific mechanistic roles of these biomarkers in endothelial phenotypic transformation.

This study has some limitations. First, the sample was derived from cells adherent to the balloon and embolic protection device during carotid artery stenting and was unavoidably mixed with blood cells. Using plaque specimens from carotid endarterectomy would likely offer greater cellular homogeneity. Second, the analysis is largely based on single-cell data, which has a limited sample size, necessitating larger cohorts for validation. Third, the findings are primarily based on bioinformatic methods; future research should incorporate animal experiments to confirm the functional roles of the identified biomarkers and provide a stronger foundation for targeted therapeutic development.

## Conclusion

KLF2, JUNB, FOS, HSPA1A, DUSP1, JUND, and ZFP36 were identified as biomarkers in this study, offering a novel perspective for elucidating the molecular mechanisms underlying ulcerative morphology in CAS.

## Non-standard Abbreviations and Acronyms

CAS: Carotid Atherosclerotic Stenosis
scRNA-seq: Single-cell RNA Sequencing
QC: Quality Control
HVGs: Highly Variable Genes
PCs: Principal Components
t-SNE: t-distributed Stochastic Neighbor Embedding
DEGs: Differentially Expressed Genes
GSVA: Gene Set Variation Analysis
GO: Gene Ontology
KEGG: Kyoto Encyclopedia of Genes and Genomes
PPI: Protein-Protein Interaction
MCODE: Molecular Complex Detection
GGI: Gene-Gene Interaction
TF: Transcription Factor
RBP: RNA-binding Protein

## Acknowledgments

The authors would like to thank Jia Yi Yin for their valuable contributions to this study, including critical manuscript review.

## Supplemental Material

Supplementary Figure 1-2

Supplementary Tables 1–8

---

**Supplementary Figure 1.** Violin plots showing the distribution of nFeature_RNA, nCount_RNA, and percent.mt before quality control (A) and after quality control (B). Highly variable gene selection (C) (Note: The y-axis shows standardized variance— greater height indicates higher gene variability. The x-axis displays mean gene expression levels. Red highlights highly variable genes, with labels showing the top 10 most variable genes. PCA analysis of the single-cell dataset (D) (Colors represent different experimental groups).

**Supplementary Figure 2.** mRNA-TF regulatory network (A) (Light purple nodes represent transcription factors predicted by individual biomarkers; dark purple indicates TFs predicted by two or more biomarkers; red nodes denote the biomarkers themselves). mRNA-RBP interaction network (B) (Red nodes mark biomarkers, orange nodes represent predicted RNA-binding proteins (RBPs), and connecting edges indicate molecular interactions).

